# Redox-driven control of Yuh1/UCHL3 impacts mitochondrial health via NEDD8/Rub1 pathway

**DOI:** 10.1101/2024.05.20.594945

**Authors:** Soha Issa, Yuval Klein, Eden Berda, Shahaf Saad, Dana Harshuk-Shabso, Abhishek Sinha, Yehonatan Sharaabi, Moran Benhar, Elah Pick

**Affiliations:** Department of Human Biology, University of Haifa, Haifa 31905, Israel; Department of Evolutionary and Environmental Biology, University of Haifa, Haifa 3190500, Israel; Department of Biology and Environment, University of Haifa at Oranim, Tivon 3600600, Israel; Department of Biochemistry, Rappaport Faculty of Medicine, Technion–Israel Institute of Technology, Haifa 3109601, Israel

**Keywords:** Reactive oxygen species, NEDD8/Rub1, ubiquitin, Cullin neddylation, Yuh1, Redox regulation

## Abstract

The ubiquitin-like protein NEDD8/Rub1 undergoes processing by the enzyme Yuh1/UCHL3 to become functional. While the processed NEDD8/Rub1 modifies Cullin-RING E3 ligases (CRLs) among all studied organisms, its role in facilitating CRL-based substrate degradation is absent in *S. cerevisiae*. This prompts questions about NEDD8/Rub1 functionality if it does not activate CRLs universally. Previous studies revealed that increased production of reactive oxygen species (ROS) during the glycolysis to mitochondrial respiration transition inhibits cullin NEDDylation in *S. cerevisiae*, yet the specific affected enzymes remain unidentified. Here, we investigate how redox changes affect Yuh1 activity, revealing a thiol-based redox switch modulating its catalytic function in response to ROS. Temporal inactivation of Yuh1 fine-tunes NEDD8/Rub1 mature and precursor species, both crucial for maintaining mitochondrial integrity and enhancing oxidative stress resilience. These findings unveil a novel role for Rub1/NEDD8 beyond CRL activation, linking redox signaling to NEDD8/Rub1 pathways.

**Graphical abstract:** 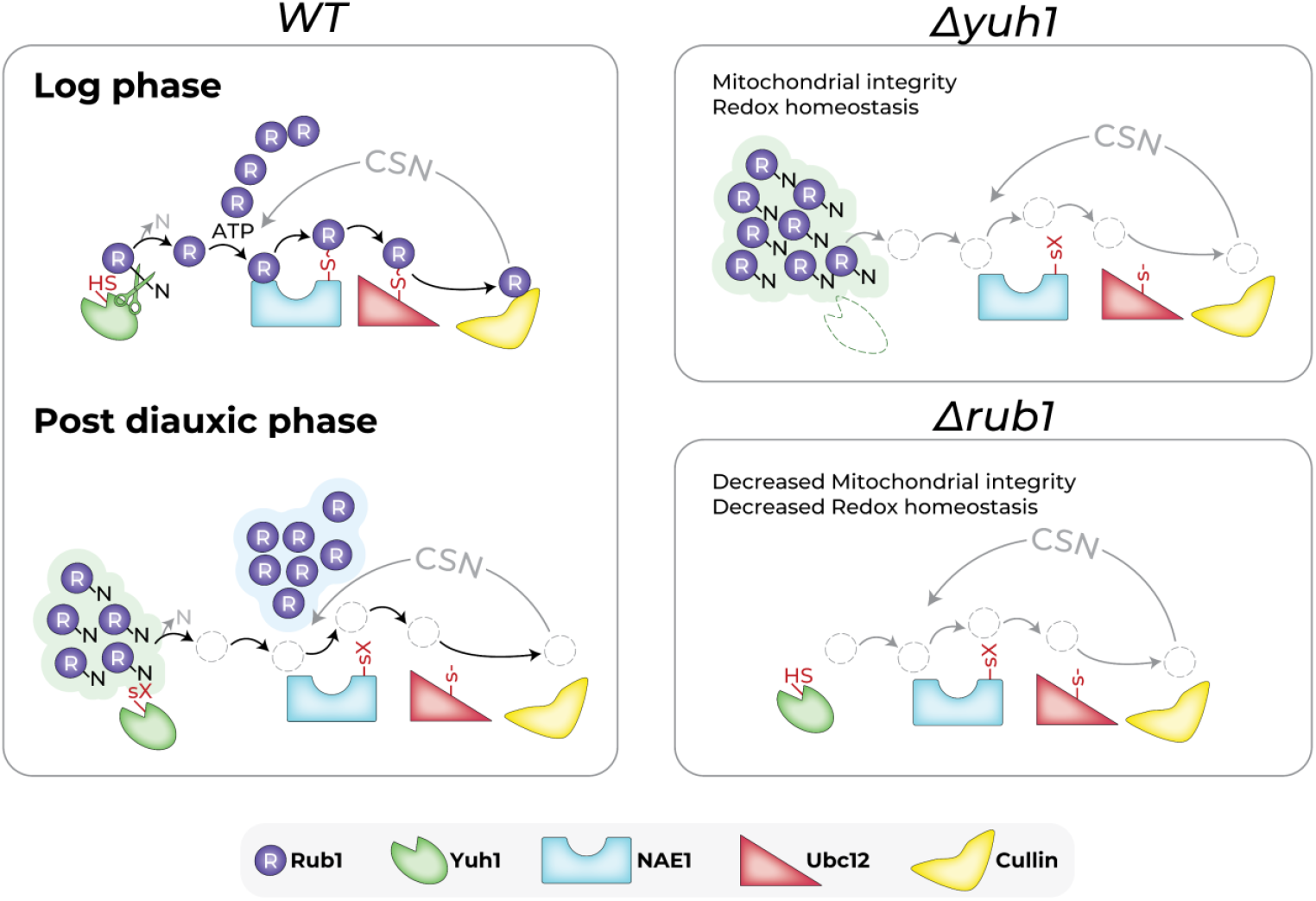

## Introduction (without heading)

Cysteine (Cys) thiols (R-SH) are among the most reactive amino acid side chains, able to function as sensors and transducers of redox status ^1^. Thiols linked to redox-sensitive cysteines, with lower pKa values, undergo deprotonation at physiological pH, forming reactive thiolate anions (R-S-). Compared to thiols, thiolates are more nucleophilic, making them more susceptible to oxidation ^2^. Redox-sensitive thiols play crucial roles in key cellular mechanisms, including the post-translational modification (PTM) of proteins by ubiquitin (Ub) and Ub-like modifiers (Ubls) ^3^. The attachment of Ubls to target proteins involves dedicated enzymatic cascades. Each cascade begins with activation of the Ubl by an ATP-consuming E1 enzyme, forming an Ubl∼E1 thioester between a C-terminal diglycine (GG) motif of the Ubl and an active-site thiol in the enzyme. The next step entails the transfer of Ubl from E1 to the catalytic thiol of an E2-conjugating enzyme, resulting in the formation of an E2∼Ubl thioester bond. Finally, the Ubl is transferred to amino groups on target proteins in a step mediated by a substrate specific E3 ligase ^4^. Despite the presence of thiol in active sites, it is assumed that Ubl cascades are not targeted by oxidation due to a relatively high pKa of E2 active sites ^5^. Still, ROS target few Ubl enzymes, suggesting that production of thiolates following the transfer of Ubl could be susceptible to oxidation ^6,7^.

NEDD8 is the closest paralog of Ub in terms of sequence and structure ^8^. Important targets of NEDD8 are “cullins” (Cul1-3, 4a/b, 5, and 7 in humans; Cdc53/yCul1, yCul3, and Rtt101/yCul4 in *S. cerevisiae*), scaffolding component of **C**ullin-**R**ING E3 **L**igases (CRLs), a large class of modular Ub-E3 ligases ^9,10^. NEDD8 attachment (NEDDylation) activates CRLs and enhances their ability to ubiquitinate specific substrates for proteasomal degradation, a critical mechanism for proper cellular functions ^11^. NEDD8 is translated as a precursor protein that undergoes maturation by proteases to reveal its C-terminal di-glycine (GG) motif, essential for its function. In humans, two proteases, UCHL3/Yuh1 and NEDP1 are responsible for NEDD8 maturation, whereas only Yuh1 is present in *S. cerevisiae* ^*12*,*13*^. Following its maturation, NEDD8 (a.k.a Rub1 in *S. cerevisiae*) is primed for cullin modification via a highly conserved enzymatic pathway. This pathway initiates with the NAE1 heterodimeric Rub1 E1 activating enzyme, (consisting in *S. cerevisiae* of Ula1 and Uba3), followed by the E2 conjugating enzyme (Ubc12 in *S. cerevisiae*), and culminates with the RING E3 subunit of CRLs (Hrt1/Rbx1 in *S. cerevisiae*). The hydrolysis of NEDD8/Rub1 from cullins is exclusively carried out by the COP9 signalosome (CSN), an evolutionarily conserved multi-subunit protease that regulates the turnover of hundreds of CRL substrates ^14,15^. While cullins are modified by NEDD8/Rub1 in all studied organisms, this modification does not activate CRLs in ascomycetes such as *S. cerevisiae*.

In a previous study, we revealed that the NEDDylation cascade of *S. cerevisiae* is sensitive to the increased mitochondrial ROS induced at the diauxic shift (DS), a physiological transition from anaerobic glycolysis to mitochondrial respiration accompanied by high ROS production ^16^. As a result, both Ubc12∼Rub1 thioester forms and cullin-Rub1 conjugates decreased. Comparable results were found in human cells upon exposure to H_2_O_2_^17^. So far, it remains uncertain whether all enzymes involved in the NEDDylation cascade are subject to thiol-based redox control.

In this study, we used budding yeast as a model organism to investigate the properties of the NEDDylation cascade under conditions of ROS accumulation. Unlike most organisms where glycolysis is coupled with oxidative respiration, budding yeast exhibits decoupled aerobic fermentation and oxidative respiratory metabolic phases. This system enabled us to identify Rub1 as a pivotal factor crucial for cellular health under conditions of ROS accumulation, with the Yuh1 protease fine-tuning the balance between matured (Rub1-GG) and precursor (Rub1-GGN) forms. These forms are both involved in sensing redox and promoting cellular integrity and survival.

## Results (divided by topical subheadings, up to 6 items -figures and/or tables)

### The inhibitory effect of metabolic ROS production and H_2_O_2_ treatment on cullin NEDDylation

We evaluated alterations in Ubc12∼Rub1 thioester formation during the DS employing tris(2-carboxyethyl) phosphine (TCEP), used to reduce disulfide bonds and sulfenic acid but maintain the Ubc12∼Rub1 thioester forms ^18^. Indeed, after 8 hours of growth, near the DS, Ubc12∼Rub1 thioester forms vanished. These observations were in line with the significant decrease in yCul1 NEDDylation status (**Fig. 1A**) ^16^. Similar effects were observed under conditions of glucose depletion, characterizing the DS (**Fig. 1B**). It is noteworthy that both the DS and glucose starvation are associated with the accumulation of ROS such as H_2_O_2_, an inevitable byproduct of electron leakage from the respiratory chains to molecular oxygen during oxidative phosphorylation in the mitochondria (**Fig. 1D**). To support this, we compared ROS production during various yeast growth phases and following a 10-minute treatment of logarithmic yeast cells with H_2_O_2_ (1.2 and 2.4 mM). The data revealed that ROS levels after treating logarithmic cells with 1.2 mM H_2_O_2_ for 10 minutes were comparable to those occurring during natural oxidation at the DS or glucose starvation. In line with this, the treatment of yeast cells with 1.2 mM H_2_O_2_ resulted in the loss of thioester formations within 10 minutes (**Fig. 1C, D**). These observations collectively suggest that thioester formation and cullin NEDDylation status are sensitive to ROS levels.

**Figure 1.**
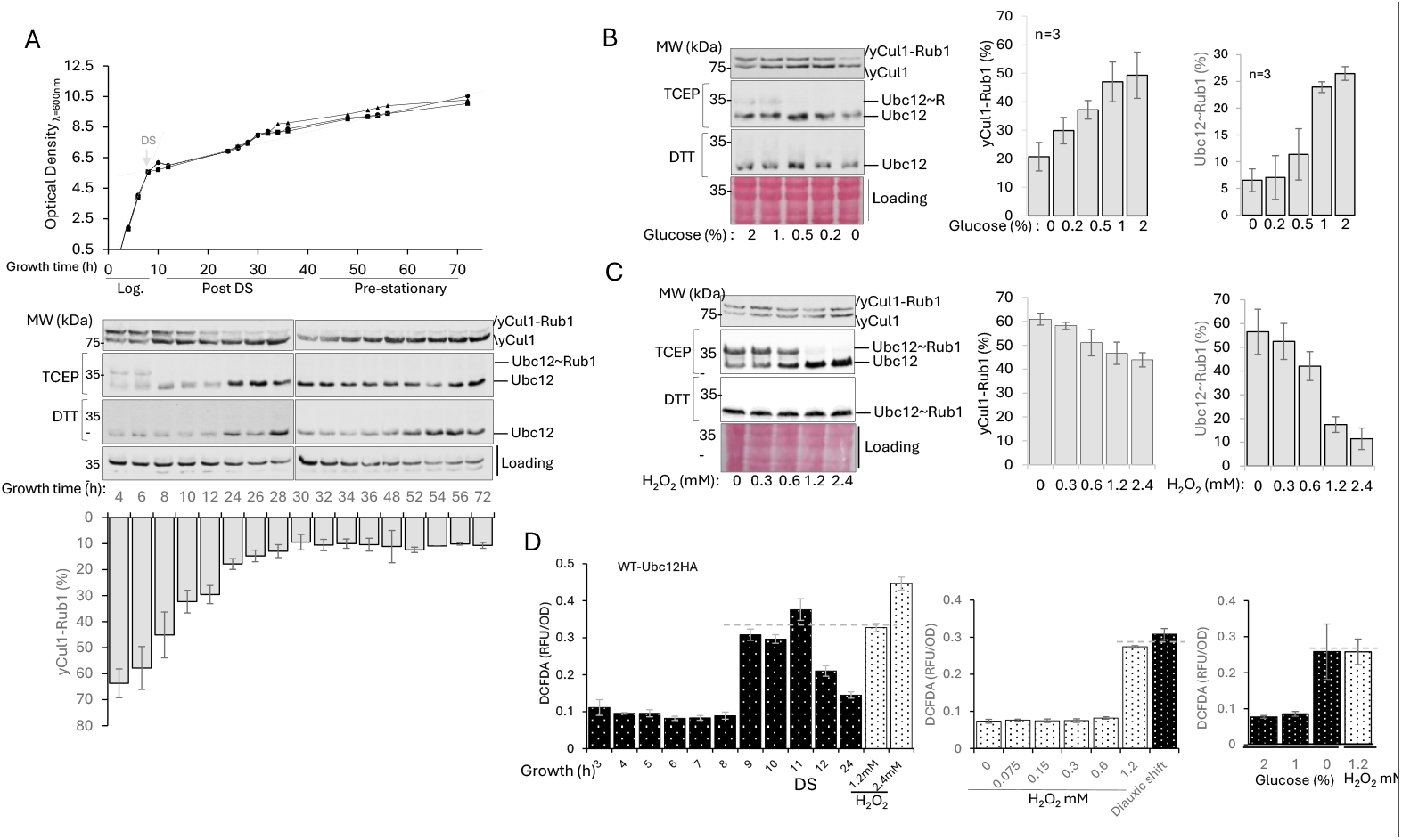
The oxidative state of *S. cerevisiae* cells upon natural and conditional induction of ROS. (A) Overnight grown wild-type (WT) cell cultures harboring genome-engineered *UBC12* ORF, C terminally tagged with 3HA, were diluted with 4% glucose YPAD to 0.5 OD600 and incubated at 28°C for 72 hours. Culture densities were measured at the indicated time, and the diauxic shift (DS) is depicted (top). Total cells extracts were collected for immunoblotting (middle) to evaluate Rub1 thioesters and conjugates while DTT was used to monitor the reduced forms of Ubc12 and yCul1. Immunoblots were quantified using ImageJ software (n≥3) (bottom). (B) Logarithmic phase cultures were grown in synthetic defined medium supplemented with 0-2% glucose and incubated at 28°C for 2 hours. (C) Logarithmic phase cultures were supplemented with 0-2.4 mM H_2_O_2_ for 10 minutes. (B, C) Cultures were harvested for immunoblotting to assess yCul1 NEDDylation status, and Ubc12∼Rub1 thioester formation. Quantification was done using ImageJ software (n≥3, B, C, right). (D) The accumulation of reactive oxygen species (ROS) was monitored using 2′,7′-dichlorofluorescein diacetate (DCFDA) in growing cells over time (left), in logarithmic cells treated with H_2_O_2_ for 10 minutes (middle), or upon glucose starvation (right). ROS levels were quantified as “relative fluorescence units” (RFU) per cell density. Notably, no significant differences (p > 0.05) in ROS levels were observed, measured either at the diauxic shift, in glucose-free medium, or upon treatment with 1.2 mM H_2_O_2_, as determined by a t-test.

### Rapid depletion of Ubc12∼Rub1 thioester forms in response to H_2_O_2_

We assessed the time-dependent formation of Ubc12∼Rub1 thioesters post-H_2_O_2_ treatment, uncovering their rapid (within 10 minutes) decrease at H_2_O_2_ concentrations greater than 0.3 mM, with thioester forms beginning to recover after 30 minutes and fully recovering after 2 hours (**Fig. 2A**). The observed patterns of loss and recovery in the Ubc12∼Rub1 thioester forms at 10 and 30 minutes, respectively, correlated with alterations in cullin NEDDylation status (**S Fig. 1**). Remarkably, in contrast to the effect upon glucose depletion and H_2_O_2_ exposure, treatment with the thiol-oxidizing agent diamide did not lead to decreased formation of Ubc12∼Rub1 thioesters (**Fig. 2B**). Cell exposure to diamide causes the rapid oxidation of reduced glutathione (GSH) to GSSG, thereby inducing a redox imbalance in GSH/GSSG. Conversely, exposure to H_2_O_2_ leads to an increase in intracellular peroxide levels, directly oxidizing sulfur-containing amino acids and generating OH• radicals. This disparity suggests that Ubc12∼Rub1 thioesters formation is less sensitive to GSH status. Overall, the findings lend support to one of two scenarios: either Ubc12 is a direct target of oxidation, or Ubc12∼Rub1 thioesters are decreased due to the action of an upstream ROS-sensitive enzyme, thereby inhibiting the transition of Rub1 downstream to Ubc12. Notably, H_2_O_2_ deactivate SUMO E1 and E2 enzymes by promoting the formation of a disulfide bond between their catalytic cysteines ^19^. We examined HMW forms, exceeding the size of Ubc12∼Rub1 (i.e., exceeding 35kDa). However, the results did not reveal the presence of disulfide-linked species following oxidation (**S Fig. 2A**). This suggests that the inhibition of cullin NEDDylation by ROS involves a mechanism distinct from that of SUMOylation.

**Figure 2.**
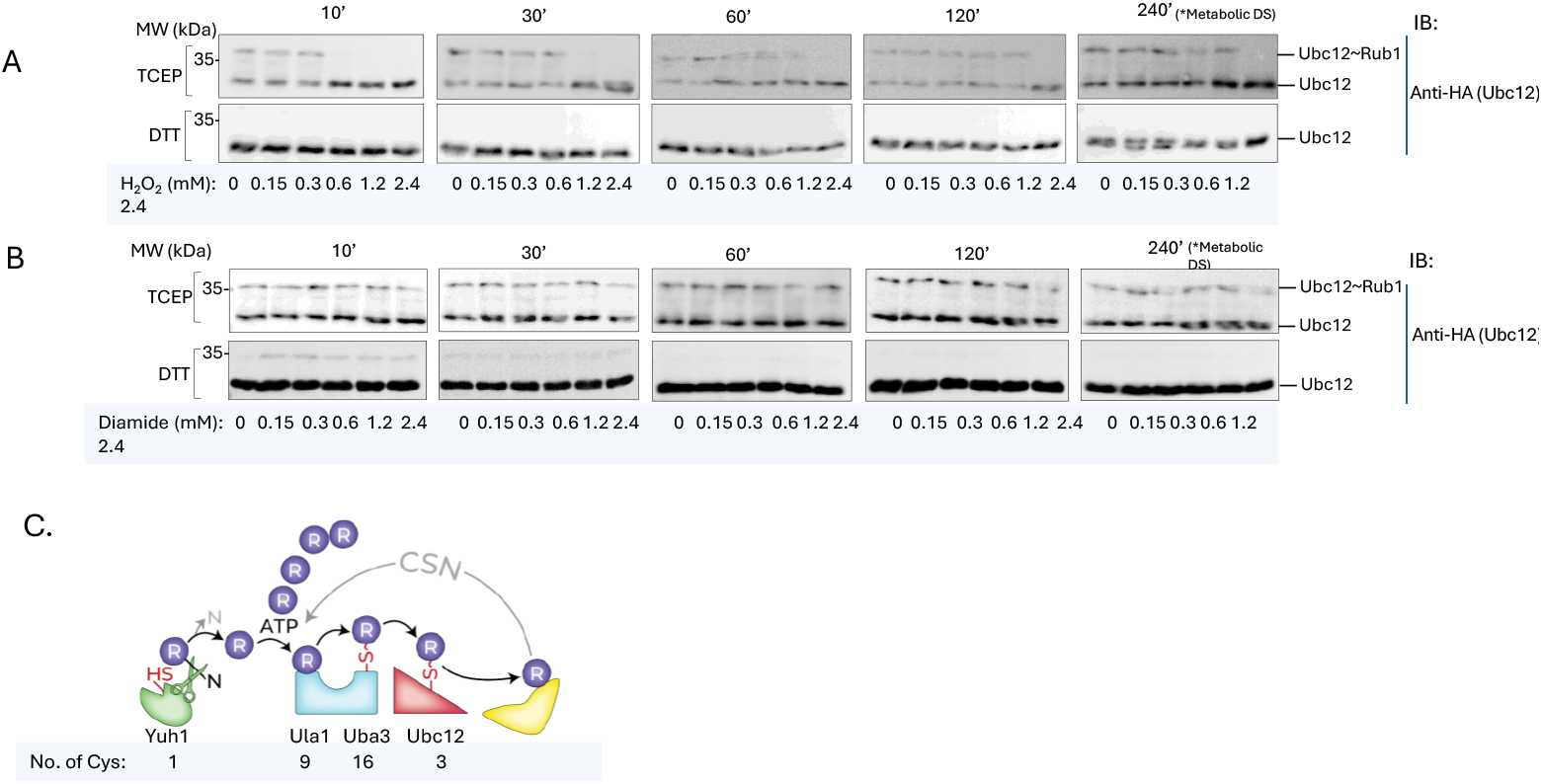
Alterations in Ubc12∼Rub1 thioester form upon H_2_O_2_ treatment. (A) Overnight cultures of wild-type (WT) cells expressing endogenous Ubc12-HA were diluted with 4% glucose YPAD to 0.5 OD600 and incubated for four hours (logarithmic phase). H_2_O_2_ (0-2.4 mM) was added directly to the growth media and incubated as indicated. Ubc12∼Rub1 thioester forms under non-reduced (TCEP) and reduced (DTT) conditions were tested by immunoblotting. (B) A similar experimental setting was employed to assess the effect of diamide on thioester formations. (C) A schematic representation of the NEDDylation cascade, starting with Rub1 maturation by Yuh1, followed by the NAE1 enzyme Ula1/Uba3 and the Ubc12 E2 enzyme, indicating the numbers of their cysteine residues.

### Inhibition of Yuh1 enzymatic activity by oxidative stress

The NEDDylation cascade, initiated by Yuh1, involves three catalytic thiols alongside additional Cys residues: a singular Cys residue in Yuh1, 25 Cys residues in NAE1, and 3 Cys residues in Ubc12 (**Fig. 2C**). To assess Yuh1 enzymatic activity in oxidative environment, we utilized a *Δyuh1* yeast strain expressing YUH1-Flag-6His under its basal promoter or an empty vector as a control ^20^. Native total cell extracts (TCEs) were prepared from transformants (**S Fig. 3A**). The fluorogenic substrate 7-amido-4-methylcoumarin (AMC), released from the carboxy terminal of Ubiquitin-AMC (Ub-AMC), was employed to measure Yuh1 catalytic activity. Our data confirmed Ub-AMC as a major substrate for Yuh1 (**Fig.3A**, ^20^). This activity was efficiently inhibited by PR619, a general Cys deubiquitinase inhibitor and by TCID, a specific human UCHL3 inhibitor, suggesting conservation of Yuh1 active site across phyla and confirming that the assay is highly sensitive to Yuh1 (**S Fig. 3B**). To verify if Yuh1 is functional in oxidized environments, we conducted experiments with various concentrations of H_2_O_2_. The results revealed a dose-dependent decrease in Yuh1 activity, with a complete inhibition, occurred at 0.15 mM of H_2_O_2_ (**Fig.3B**). These findings were further supported when a bacterially expressed Yuh1, evaluated for its ability to cleave Ni-NTA based pre-purified 6His-UBB+1 (a carboxyl terminal extended mutant of Ub), exhibited H_2_O_2_-induced inhibition. Control experiments using recombinant commercial Ulp1 and Yuh1-free TCE from E. coli BL21 corroborated the redox-sensitive nature of Yuh1 (**Fig.3C**). Yuh1 inhibition proved reversible, with full activity restoration achieved by the addition of DTT to the H_2_O_2_-treated TCEs (**Fig.3D**). Collectively, the above findings highlight Yuh1 as a redox-sensitive enzyme.

**Figure 3.**
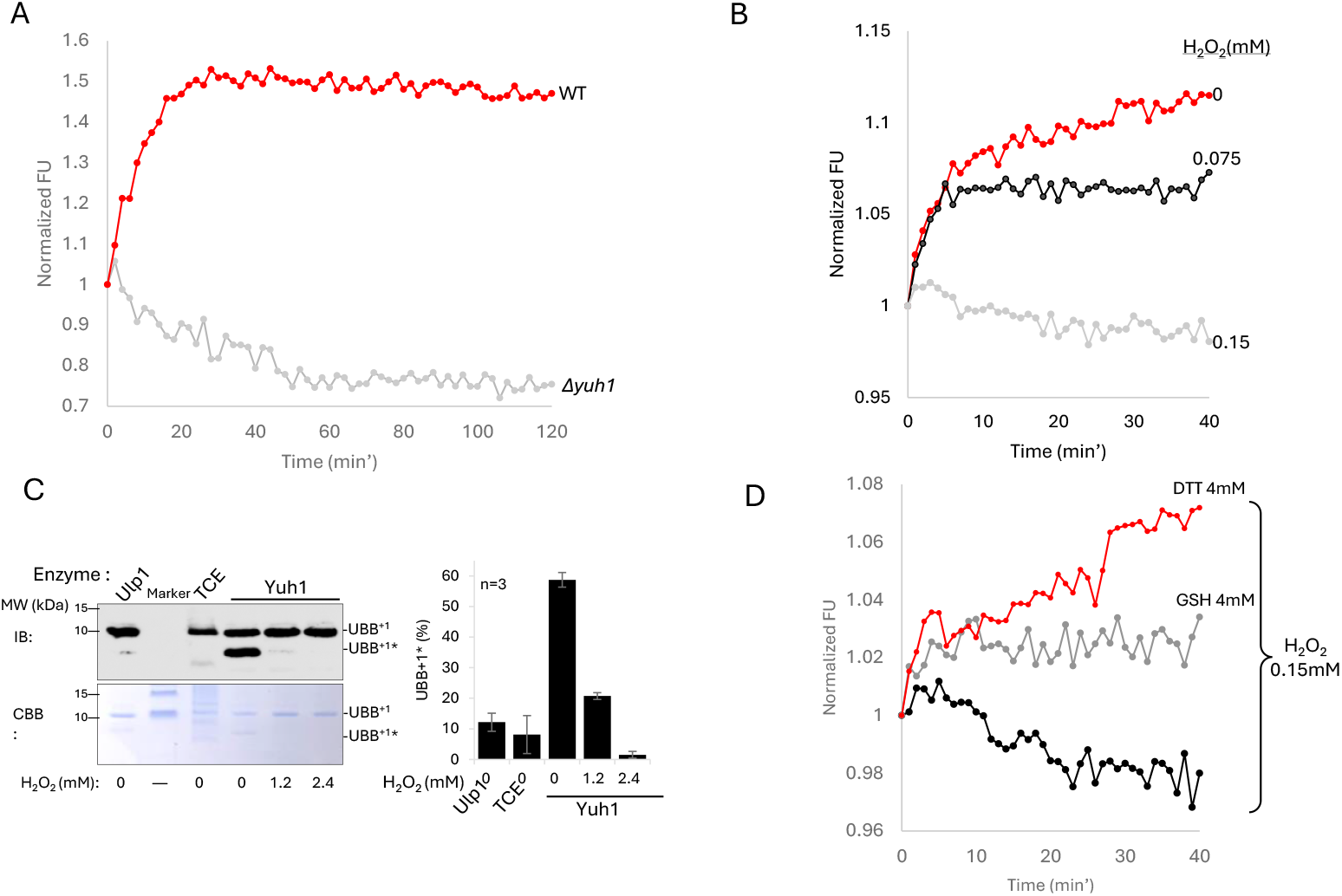
Yuh1 activity is inhibited by hydrogen peroxide. A mutant strain of *Δyuh1*, complemented by Yuh1-6His-Flag under *YUH1p* promoter (WT) or an empty plasmid (*Δyuh1*) was cultivated to the logarithmic phase at 28°C. (A) Native total cell extracts (TCE) were utilized to assess Yuh1 activity by a plate reader, measured as the cleavage ability of the amide bond between Gly76 of ubiquitin and 7-amido-4-methylcoumarin (AMC). The plot illustrates the relative fluorescence units (RFU), normalized at each time point relative to the first read. Representative result n≥3. (B) The ability of Yuh1 to cleave Ub-AMC was measured in WT lysates treated with various concentrations of H_2_O_2_ (0.075,0.15 mM) for 10 minutes, without (B), or with (D) the addition of DTT and glutathione (GSH) antioxidants. Representative result n≥3. (C) Yuh1 activity in full lysates of yeast WT was measured in the presence of H_2_O_2_ (1.2 and 2.4 mM) for 10 minutes, using the recombinant protein UBB^+1^ as a substrate. Results were visualized by Coomassie brilliant blue (CBB) staining and immunoblotting (IB) (left). The data from three repetitions were analyzed using ImageJ (right). Recombinant Ulp1 SUMO hydrolyzing enzyme and a total cell extract (TCE) of BL21 bacteria were used as controls for enzymatic activity. Representative result n≥3.

### Targeting a reversibly modified redox-sensitive cysteine in Yuh1

The decrease in Yuh1 activity in an oxidative environment may result from redox-based thiol modification. To explore this, we initially investigated the susceptibility of one of the three thiols in Ubc12 and the single thiol in Yuh1 to oxidation using a gel-shift assay. This assay involved the use of Methoxy PEG Maleimide (mPEG-MAL), a thiol-reactive PEG derivative, which modifies free thiols, leading to size shifts detected by immunoblotting. We conducted an experiment where mPEG-MAL was introduced to extracts from cells expressing Yuh1-His-Flag and endogenous Ubc12-HA pre-treated with H_2_O_2_. The results showed a significant alteration in the sizes of Ubc12 and Ubc12∼Rub1, indicating that multiple free thiols of Ubc12 were targeted by mPEG-MAL. Importantly, this shift occurred independently of H_2_O_2_ treatment, suggesting that Ubc12 cysteines are not readily oxidized **(S Fig. 4A)**. In contrast, we found that Yuh1 was clearly and dose-dependently oxidized in response to H_2_O_2_, indicating direct oxidation of its catalytic Cys-thiol group **(Fig. 4A)**. Notably, the source of mPEG-MAL free Yuh1 does not originate from the translation of a new protein, as its quantity remained constant even with the addition of cycloheximide. This contrasts with Cdc4, a short-lived protein used as a control **(S Fig. 4B)**. To explore possible reversibility of Yuh1 Cys-thiol oxidation, we employed a PEG-switch assay, starting with H_2_O_2_-induced oxidation, followed by the blocking of non-oxidized thiols using Iodoacetamide (IAM). Subsequently, oxidized cysteines were reduced by TCEP to enable alkylation by mPEG-MAL, resulting in approximately five kDa mobility shift for each free thiol in SDS-PAGE. Immunoblotting for Yuh1 revealed an H_2_O_2_ dose-dependent alkylation of the active site thiol of Yuh1 **(Fig. 4B, S 4C)**. The results clearly indicate that Yuh1 undergoes reversible modification in response to ROS, while Ubc12 remains largely unaffected. Moreover, in correlation with **Fig. 2A**, the results revealed a time-dependent decline in the PEGgylated conjugates, suggesting oxidative stress recovery over time **(Fig. 4C)**. The reversible modification observation was corroborated using the Ox-RAC method, specifically designed for capturing oxidized proteins. Like the PEG-switch assay, this method involves the capture of reduced thiols, employing thiopropyl sepharose (TPS) resin, which captures proteins containing free thiols via a disulfide exchange reaction ^21^. The captured proteins were eluted using TCEP and digested for MS analysis of cysteine-based reversible modifications. Significantly, Yuh1 but not Ubc12 was predominantly captured in the oxidant-treated samples, as confirmed through both immunoblotting and LC-MS/MS analyses (**S Fig 5, S Table 4**). Altogether, Yuh1 activity is subject to reversible inhibition by H_2_O_2_.

**Figure 4.**
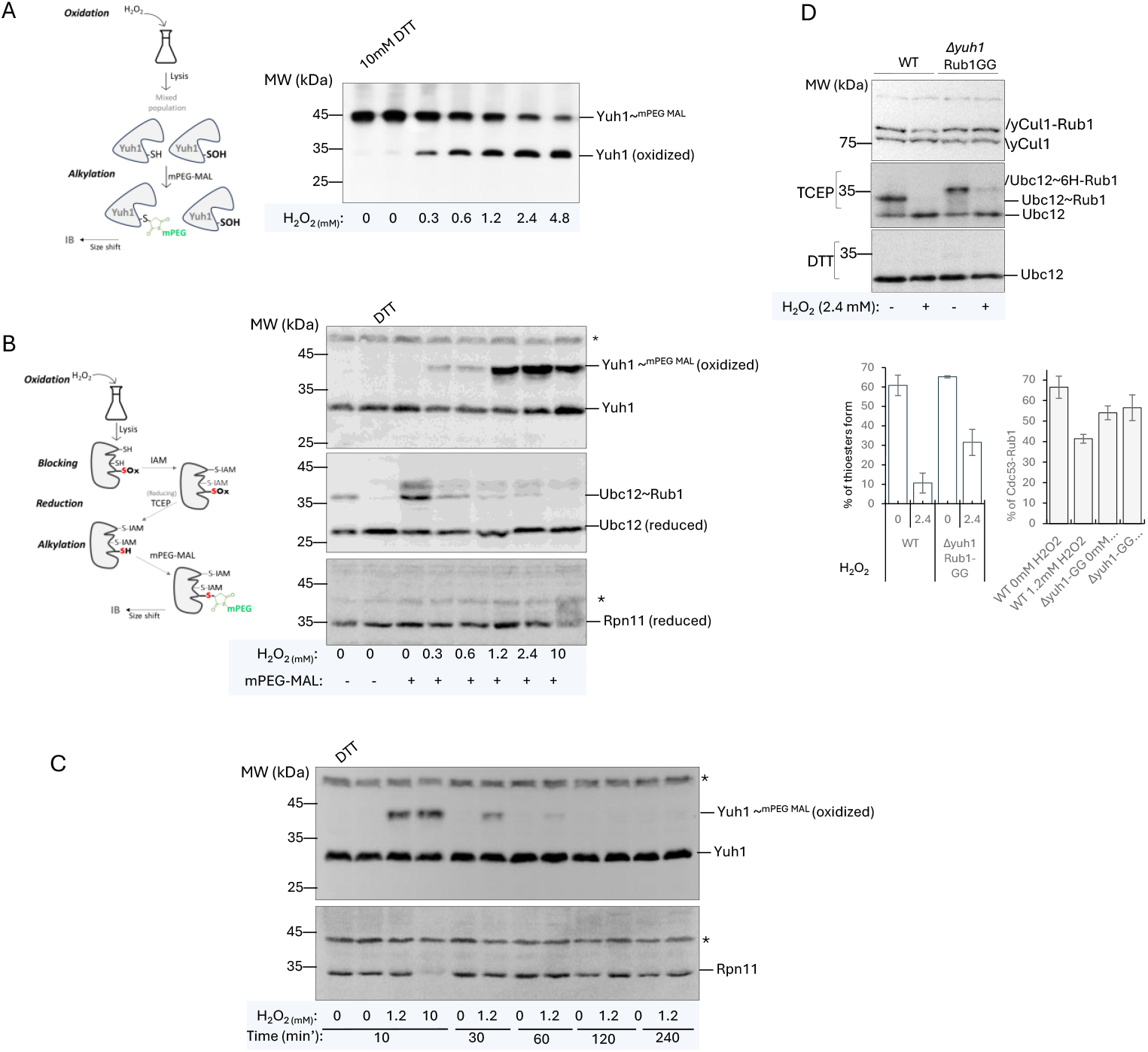
Confirming the oxidation of Yuh1 Cys by gel-shift PEGylation. (A) Left - Schematic representation of Yuh1 PEGylation assay. Right - Overnight grown *Δyuh1* mutant cells co-expressing endogenous Ubc12-HA and ectopic Yuh1-His-Flag were diluted with 4% glucose YPAD and incubated to the logarithmic phase prior to treatment with the indicated concentration of H_2_O_2_ for 10 minutes or 10 mM DTT for 30 minutes. Methoxypolyethylene glycol maleimide (PEG-MAL) was added to monitor changes in reduced Cys, and PEGylation of Yuh1 was examined by immunoblotting under non-reducing conditions. (B) Left - Schematic representation of Yuh1 PEGylation following the chemical blocking of “free” thiols by iodoacetamide (IAM). Right - logarithmic *Δyuh1* mutant cells expressing endogenous Ubc12-HA and ectopic Yuh1-His-Flag were treated with various concentrations of H_2_O_2_ for 10 minutes or 20mM DTT for 30 minutes. To monitor changes in reduced Cys, PEG-Maleimide was added following the addition of IAM and reduction by 10 mM TCEP to monitor changes in oxidized cysteines. (C) The same experiment was conducted with samples taken for validation overtime. The oxidation/reduction status of the various proteins in A-C (Yuh1, Ubc12 or Rpn11) was evaluated through immunoblotting. (D) Logarithmic cultures of WT Ubc12-HA and Ubc12-HA Δyuh1 mutant strains expressing the matured form of Rub1 (Rub1-GG) were treated with 1.2mM of H2O2 for 10 minutes and subsequently harvested for immunoblotting under non-reduced (top) or reduced (bottom) conditions (top). In both strains, yCul1 NEDDylation status and Ubc12 thioester formations were quantified by ImageJ software (n≥3) (bottom).

Exploring the role of Yuh1 in Rub1 activation, we aimed to unravel the ramifications of ROS-targeted inactivation of its catalytic Cys on the NEDDylation cascade. To elucidate this, we employed *Δyuh1* cells overexpressing Rub1-GG as a mechanistic approach to investigate the NEDDylation cascade in a system bypassing the need for Yuh1. **(Fig. 4D, S6)**. Our findings revealed that, compared to the WT, this strain retained a moderate quantity of Ubc12∼Rub1 thioester forms even under ROS accumulation, sufficient to sustain yCul1 modification by Rub1. This underscores the critical role of Yuh1 in maintaining cullin NEDDylation. Nevertheless, the reduction in Ubc12∼Rub1 upon H2O2 exposure in *Δyuh1* cells overexpressing Rub1-GG indicates that NAE1 is another potential redox-sensitive NEDDylation enzyme, in addition to Yuh1, operating upstream of Ubc12. These findings prompt further investigation into whether the suppression of Yuh1 activity upon accumulated ROS exerts a positive or negative impact on cellular health.

### Yuh1 is crucial for maintaining redox balance and mitochondrial health

The inhibition of Yuh1 in oxidative environments hinders the maturation of newly synthesized Rub1 molecules. However, even in the presence of Yuh1 inhibition under conditions of oxidative stress, the CSN complex remains active. CSN activity releases a pool of matured Rub1 molecules recycled from cullins, bypassing the need for Rub1 maturation via Yuh1 (**Fig 2C, 4D**) ^22^.

In *S cerevisiae* the absence of Rub1 (*Δrub1*) typically results in mild mitochondrial defects ^16^. These phenotypes were reversed upon the expression of UbK0,R72T, a lysine-less Ub variant designed to mimic Rub1 ^23^. In line with this observation, *Δcsn5* mutant strain, characterized by minimal levels of free Rub1, demonstrated an accumulation of intrinsic ROS ^22^. Contrariwise, mutants such as *Δuba3, Δubc12*, and *cdc53k760/R*, accumulate the matured Rub1-GG and exhibit lower ROS levels **(S Fig. 7 A, B)**. This prompts further inquiry into whether inhibiting Yuh1 to prevent Rub1-GGN to Rub1-GG maturation alters mitochondrial and redox-based properties of cells. Consequently, we explored the necessity of the Rub1-GGN precursor during oxidative stress. Interestingly, endogenous ROS levels in Δ*yuh1* mutant cells exceed those in the WT strain (**S Fig. 7C**). However, overexpression of either mature Rub1-GG or its precursor Rub1-GGN in Δ*yuh1* mutant cells effectively mitigates this increase, indicating that full maturation of Rub1 is not necessary for its protective role against oxidative stress (**Fig. 5A, S 7C, D**). Viability assays under acute oxidative stress conditions (20mM for one hour) revealed that *Δrub1* exhibits lower viability than *Δyuh1*, particularly under conditions favoring mitochondrial respiration with glycerol as the carbon source (YPGly). This suggests differential mitochondrial utilization between the strains and implicates a role for the Rub1-GGN precursor naturally harbored within *Δyuh1* (**Fig 5B, S7E**).

**Figure 5.**
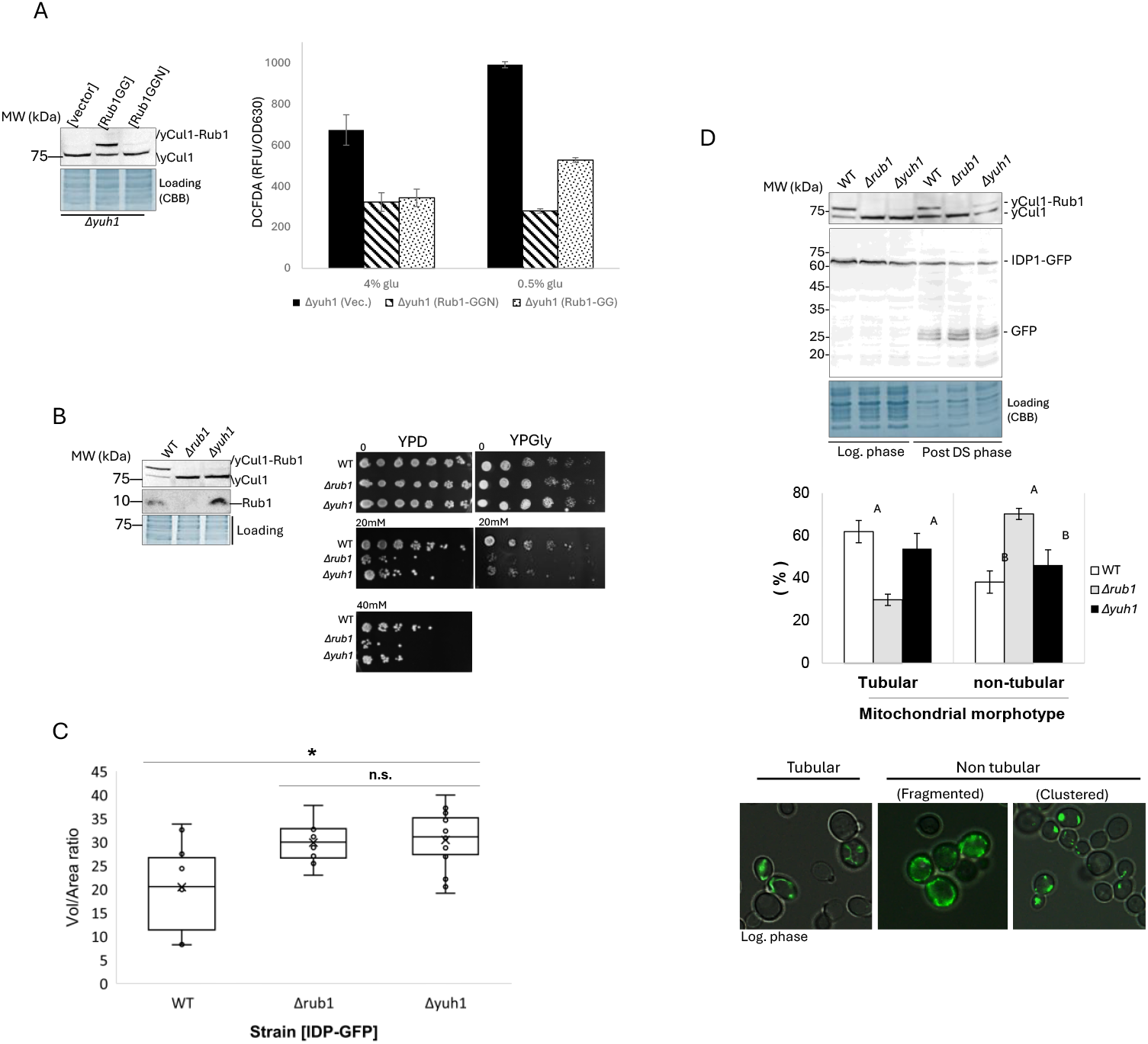
Impact of Yuh1 activity on cellular health. (A) Ectopic expression of the Rub1-GGN precursor or the matured form Rub1-GG in *Δyuh1* cells. Left: Immunoblotting for yCul1 confirmed that only Rub1-GG modifies it. Right: Logarithmic culture was grown in SC medium supplemented with 4% or 0.5% glucose for 2 hours and pretreated with DCFDA. Fluorescence signal was measured as described in Fig 1D. (B) Logarithmic WT, *Δyuh1* and *Δrub1* mutant cells were treated with 20mM, or 40mM of H_2_O_2_ for 1 hour to induce acute oxidation (right). Recovery was evaluated by seeding treated cultures onto YPAD or YPGly plates using a serial drop assay. (C) Mitochondrial volume to cell area ratio of the WT compared to *Δrub1* and *Δyuh1* S. cerevisiae strains based on their confocal microscopy images. Asterisks indicate significant differences between strains (p<0.05) and n.s. indicate a non-significant difference (p>0.05) by Bonferroni post-hoc tests. (D) Mitochondrial morphotypes (%) of the three studied *S. cerevisiae* strains. Top - WT, *Δyuh1* and *Δrub1* Cells ectopically expressing IDP1-GFP were subjected to Immunoblotting for yCul1 and GFP for IDP1 at the indicated growth phases. Bottom - Strains were evaluated for mitochondrial morphotypes in the total cell population, and the average count from five different areas was calculated. Different letters indicate significant differences between groups based on two-way ANOVA (p<0.05).

Mitochondria, recognized as significant contributors to ROS production, undergo dynamic morphological changes in response to oxidative stress ^24^. During the DS, yeast cells undergo a surge in mitochondrial biogenesis, accompanied by an increase in mitochondrial volume respiratory conditions ^25,26^. Our assessment of mitochondrial volume to cell area ratio indicates that the WT strain was significantly different from that of the mutant strains (One-way ANOVA: F_2,29_ = 6.35, p = 0.005). The results point out an increase of approximately 32% in mitochondrial volume to cell area ratio in *Δyuh1* and *Δrub1* mutants compared to WT strains. The lowest ratio was observed in WT, with a mean ratio of 20.41 ± 3.03. In comparison, both *Δrub1* and *Δyuh1* had significantly larger ratios, with means of 29.88 ± 3.03 and 30.41 ± 1.69, respectively. Bonferroni post-hoc test revealed a significant difference in the WT ratio compared to *Δrub1* and *Δyuh1* (p = 0.03 and p = 0.007, respectively). Interestingly, the standard errors of *Δrub1* and *Δyuh1* were nearly identical, albeit 56% lower than in the WT population. The more uniform mitochondrial volume in these mutants, with reduced fluctuations, may indicate synchronized cellular responses to pressure constraints **(Fig. 5C, D)**. Nevertheless, the findings showed no significant difference between the *Δrub1* and *Δyuh1* mutants (p>0.05), implying that both expression and maturation of Rub1 play crucial roles in maintaining mitochondrial volume.

Another aspect demonstrating the dynamic nature of yeast mitochondria is the formation of tubular-like filamentous morphotypes, resulting from a delicate balance between fission and fusion events. During the logarithmic phase, yeast cells predominantly feature tubular mitochondria when cultured in glucose containing YPAD medium. However, upon glucose depletion at the DS, mitochondria undergo fragmentation in a Dnm1-dependent manner, transitioning to a more clustered morphology typical of the stationary phase ^27-29^. Oxidative stress similarly triggers excessive mitochondrial fragmentation and clustering ^30^. To evaluate mitochondrial morphology in *Δrub1* and *Δyuh1* mutants, we expressed the mitochondrial NADP-specific isocitrate dehydrogenase (IDP1) tagged with GFP in cells. This method has been previously utilized to evaluate mitochondrial dynamics via microscopy and observe mitophagy in stationary phase cells, confirmed by GFP release observed through immunoblotting ^31,32^. We confirmed that unlike in the 3-day-old culture, IDP1-GFP remains uncleaved in logarithmic cells, effectively labeling the mitochondria (**Fig. 5D top**). Our observations delineated distinct mitochondrial phenotypes during the logarithmic phase: while the majority of WT and *Δyuh1* mutant cells exhibited the expected tubular mitochondria morphotype, *Δrub1* mutant cells displayed pronounced mitochondrial fragmentation (**Fig. 5D bottom**). Indeed, the two-way ANOVA revealed a significant interaction between the strain and its mitochondrial morphotype composition (Two-way ANOVA: R2adj=0.85, F2,20 = 68.85, p < 0.001). The WT strain had the highest percentage of tubular mitochondria at 61.86 ± 3.02 %, whereas this pecentage was 14% lower in the *Δyuh1* mutant strain (53.14 ± 4.15 %). Interestingly, the *Δrub1* mutant strain had the lowest portion of tubular mitochondria, 50% lower than the WT (30.39 ± 1.31 %). These findings suggest that *RUB1* expression mitigates mitochondrial fragmentation during the fermentative phase of yeast. Furthermore, the data highlight a novel role for the endogenous levels of Rub1-GGN accumulated in *Δyuh1* in maintaining mitochondrial tubular morphology even under the heightened endogenous ROS levels observed in this mutant (**S Fig. 7C**), while also contributing to cellular health in response to acute ROS levels (**Fig 5B, S7E**).

## Discussion

Here, we sought to decipher the mechanism by which cullin-NEDDylation—a process mediated by thiol-based enzymes—is inhibited by ROS. Several pieces of evidence led us to assume that Ubc12 is the oxidation target causing the dramatic loss of cullin NEDDylation under oxidizing conditions. Indeed, a previous study that utilized nitrosothiol capture and enrichment techniques with a thioredoxin trap mutant (Trx1C35/S) revealed the capture of oxidized human Ubc12 in a large-scale oxidized protein screen ^33^. Another study employed biotinylated iodoacetamide (BIAM) and fluorescein-5-maleimide (F5M), revealed the redox sensitivity of the reactive cysteine in the mammalian Ubc12 ^17^. However, unlike the mammalian Ubc12/UBE2M, *S. cerevisiae* Ubc12 does not appear susceptible to direct cysteine oxidation. Efficient oxidative modification of cysteine necessitates its thiolate anion form, which is less prevalent at physiological pH. The surrounding amino acids of the catalytic cysteine in yeast Ubc12 differ from those in human Ubc12/UBE2M, resulting in a lower pKa of the active site (6.73 in yeast, 4.35 in human), potentially contributing to the differential susceptibility to oxidation between yeast and human Ubc12 enzymes. Our study also highlights the importance of considering reducing reagents in experimental setups involving catalytic cysteine-dependent processes like Ubl cascades. DTT, for instance, can erroneously suggest reversible oxidation by reducing the catalytic thioester within Ubc12, while TCEP avoids such misinterpretations by not affecting thioester forms.

Our study features the role of Yuh1 in modulating the balance between the Rub1-GGN precursor and the Rub1-GG mature form. The sole thiol group of Yuh1 is subject to reversible redox-based inhibition, a phenomenon validated through both Ub-AMC hydrolysis activity assays and thiol capturing assays. Inhibition of Yuh1 leads to a moderate increase in UBC12∼Rub1 thioester forms, evidenced by the persistence of matured Rub1 in Yuh1-deficient cells. This observation raises a pivotal question: What is the physiological advantage of limiting the pool of mature Rub1 through Yuh1 inhibition? Our data reveal that both Rub1-GG and Rub1-GGN effectively mitigate intrinsic ROS levels in *ΔYuh1* cells. Intriguingly, under low-glucose conditions, which simulate the diauxic shift, Rub1-GGN more effectively counteracts accumulated ROS compared to Rub1-GG, underscoring its potential importance during the oxidative respiration phase. Moreover, cell viability assays following acute ROS exposure demonstrate enhanced survival in *ΔYuh1* cells, compared with *ΔRub1*. This discrepancy is particularly evident when ROS exposed cells are cultivated on glycerol, a carbon source known to enhance mitochondrial respiration. Consistent with this, the mitochondria of logarithmic *ΔRub1* cells manifest distinct characteristics compared to *ΔYuh1* and WT strains, by displaying reduced tubularity and increased clustering, a phenotype associated with post-logarithmic cell phases and cells subjected to oxidative stress ^29,30^. These observations collectively suggest that the Rub1-GGN precursor plays a crucial role in cellular survival and in maintaining mitochondrial function and morphology, especially following exposure to acute ROS. However, the molecular mechanism underlying these physiological findings remains elusive A clue may lie in previous studies in mammalian cells, which have shown that the generation of unanchored NEDD8 trimers can interact non-covalently with PARP1, attenuating its activation and thereby delaying cell death induced by PARP1 overactivation during oxidative stress^34^. The unanchoring of NEDD8 might be attributed to the blocked C-terminal domain of Rub1-GGN, rendering it incapable of modifying client proteins but allowing it to serve as a substrate for the other two linked Rub1 molecules at its Lys residues. Although *S. cerevisiae* lacks a PARP1 orthologue, it possesses a gene known as PMS1 (Post-Meiotic Segregation Increased 1) that shares some functional similarities with PARP1 by playing a role in DNA mismatch repair and maintaining genomic stability, analogous to the DNA repair functions of PARP1 ^35^. Interestingly, both Ubc12 and Yuh1 show positive and negative genetic interactions with Pms1, respectively ^36,37^. This observation aligns with our findings. Further studies could evaluate whether the inhibition of Yuh1 and accumulation of Rub1-GGN modulate a redox signal through non-covalent interactions with PMS1, similar to the relationship between PARP1 and NEDD8 trimers.

Our study also emphasizes the critical role of the matured modifier, Rub1-GG, a player at the interplay between NEDDylation and ubiquitination pathways, in countering ROS. Diminished Yuh1/UCHL3 activity, coupled with the possible inhibition of NAE1, prevented the formation of Ubc12∼Rub1 thioesters in an all or none manner. This in higher organisms may compromise CRLs function, altering the ubiquitination of specific proteins and thereby influencing cellular adaptation to environmental challenges and the turnover of key cellular components. Indeed, mammalian Cul1 inactivity stabilizes the lung epithelial sodium channel (ENaC), crucial for alveolar salt-water balance ^38^. Bacterial-induced ROS oxidatively inactivates NEDDylation, leading to reversible Cul1 NEDDylation loss. This stabilizes β-catenin and NF-κB inhibitor IκB-α, impeding their recruitment to CRL1/SCF SR βTrCP ^17,39^. Redox signaling modulates turnover of central redox regulators and CRL substrates, impacting HIF-1α, NRF2, and FNIP1 during hypoxia, oxidative stress, or reducing environments in mammals ^40-42^. These players also regulate key metabolic pathways: HIF-1α boosts glycolysis, NRF2 promotes mitochondrial biogenesis, and FNIP1 fine-tunes mitochondrial respiration ^42^. Under non-oxidized conditions, all these substrates degrade, but during oxidative stress, their degradation is inhibited, ensuring cellular protection.

The involvement of Yuh1 in both NEDDylation and Ub maturation adds complexity to its regulation by ROS and its impact on cellular processes. Sensitivity of Yuh1 to ROS could have a dual effect on cellular stress responses, potentially reducing the pool of matured Ub available for conjugation to target proteins. This effect is exacerbated under oxidative stress, where there is already a decrease in the pool of free Ub due to the massive formation of K63-linked chains ^43^. Deficiencies in ubiquitin maturation might compromise the specificity and efficiency of ubiquitination reactions. This aligns with observations that oxidative stress can enhance activity of the 20S proteasome, potentially reducing the need for substrates ubiquitination ^44^.

In summary, our study highlights Yuh1 as a central regulator of Rub1, governing cellular redox balance and mitochondrial health. Several novel roles for NEDD8/Rub1 beyond its function in cullin modification have emerged across various organisms ^45^. Despite the yeast genome being relatively small and the non-essentiality of Rub1 for yeast viability, additional functions for this polypeptide remain elusive. The reversible modulation of Yuh1 activity by ROS underscores its importance, offering valuable insights into the complex interplay between redox signaling and protein modification pathways. Further investigations will uncover the precise molecular mechanisms through which Rub1 precursor maintains cellular balance under conditions of increased ROS.

## Methods

### 1. Yeast strains, media, and growth conditions

*Saccharomyces cerevisiae* cells were cultured under standard conditions at 28°C in rich YPAD medium (1% yeast extract, 1% peptone, 0.01% adenine hemisulfate, 2% glucose), unless otherwise specified. Plasmids were maintained by cultivating transformants in selective synthetic complete (SC) medium supplemented 17% (w/v) yeast nitrogen base, 0.5% ammonium sulfate and a complete mixture of amino acids to satisfy any auxotrophic requirements commonly encountered. For all experiments, overnight starter cultures were diluted to 0.5 OD600/ml and incubated for specified durations, temperatures, or treatments, as detailed in the figure legends. For H_2_O_2_ or diamide treatments, reagents were introduced to the culture for various times and concentrations as indicated in the figure legends. Yeast strains and plasmids utilized in this study are described in **S Tables 1, 2**.

### 2. Endogenous oxidative stress measurements

Intracellular ROS levels were assessed using 2′7′-dichlorofluorescein diacetate (DCFDA), as previously described ^46^. Briefly, DCFDA was introduced to the various cultures and incubated for one hour, followed by two washes with PBS. Subsequently, samples were analyzed either via fluorescence microscopy or using a BioTek Synergy HT plate reader (excitation 485/20 nm and emission 528/20 nm). For starvation treatments, the yeast culture was transferred to a glucose-deficient medium and then incubated for an hour before the addition of DCFDA. For the H_2_O_2_ treatments, H_2_O_2_ was introduced to the culture during the final 10 minutes of the DCFDA incubation period.

### 3. Denaturative protein extraction

For immunoblotting, cells were extracted by Trichloroacetic acid (TCA) precipitation method, as previously described ^47^. Briefly, 5 OD600 cells were washed and resuspended in 20% TCA following aggressive lysis with zirconia beads (0.5mm diameter) through vortexing. The lysed cells were then transferred to a new Eppendorf tube and centrifuged at 2300 x g for 5 minutes. Pellets were neutralized using Trisma base and supplemented with Laemmli sample buffer X5 (0.5% SDS, 12% glycerol, 50 mM pH 6.8 Tris-HCl, 2% β-mercaptoethanol). The samples are then heated at 95°C for 5 minutes to denature proteins and prepare them for gel electrophoresis.

### 4. Monitoring thioester forms

To monitor Ubc12∼Rub1 thioester forms, TCA pellets were dissolved in urea sample buffer (8M urea, 250 mM pH 11 Tris, 5% glycerol, 2% SDS, and 50 mM Tris (2-carboxyethyl) phosphine hydrochloride (TCEP)), followed by warming at 37°C for 5 minutes. For the reduced forms, 50mM of DTT (dithiothreitol) is added directly to the urea sample buffer before heating at 95°C for 5 minutes. This step ensures reduction of disulfide bonds, facilitating the analysis of reduced forms.

### 5. Native extraction of Yuh1 for enzymatic activities

Cells were pelleted and resuspended in a buffer containing 20mM Tris-Hcl pH7.4, 150mM NaCl, 1mM DTT, and 0.1% phosphatase/protease inhibitors, followed by sonication (40% sonication capability, 10 sets of 10 pulses with intervals for cooling), as further described ^20^.

### 6. Preparation of the recombinant proteins for in vitro assays

The Yuh1 enzyme was expressed in pET3a-based plasmids, while the 6His-UBB+1 substrate was expressed in pQE30-based plasmids. Bacterial strains BL21 (pET3a) and M15 (pQE30) were utilized for protein expression under the control of T7 and T5 promoters, respectively. Lysates underwent Ni-NTA-based purification of the 6His-tagged proteins, followed by dialysis (cut-off – 10kDa) at 4°C using dialysis buffer (Tris-HCl pH 7.4 20 mM, NaCl 150 mM, Tween 20 0.25%, EDTA 2 mM, DTT 1 mM, and glycerol 12.5%). The solubility of recombinant proteins was confirmed by SDS-PAGE gel followed by Coomassie brilliant blue (CBB) staining or immunoblotting.

### 7. Immunoblotting

Proteins separated by both reduced and non-reduced SDS-PAGE were transferred onto 0.45μm nitrocellulose membranes. Following blocking, specific antibodies listed in **S Table 3** and mentioned in figure legends, were used for proteins recognition. Enhanced chemiluminescence (ECL) was employed for detection, with images captured using a Fusion FX 6 EDGE chemiluminescence imaging system. Quantitative analysis was performed using ImageJ v1.40f software (http://imagej.net/). The NEDDylation status of yCul1 and Ubc12∼R thioester formation was determined by comparing the relative ratio of modified yCul1 and Ubc12∼R to their respective total amounts. Results are presented as the average and standard deviation of ratios from three or more biological replicates. Statistical analyses were conducted using IBM SPSS Statistics software (version 27).

### 8. Evaluating Yuh1 enzymatic activity using Ubiquitin-AMC as a substrate

Ubiquitin-AMC (Ub-AMC) was employed to assess Yuh1 activity. Yuh1 catalyzes the hydrolysis of the amide bond between Gly76 of ubiquitin and 7-amido-4-methylcoumarin (AMC), leading to increased fluorescence (excitation 360/40 nm and emission 460/40 nm) as described ^20^. Briefly, total yeast cell extracts were serially diluted in DUB buffer (600 mM NaCl, 100 mM Tris–HCl pH 7.4, 2.5 mM EDTA, 5 mM DTT) before the addition of Ub-AMC, and readings were initiated immediately. The plot illustrates the relative fluorescence units (RFU), normalized at each time point relative to the first read, which was defined as 1. This normalization ensures comparability across readings despite potential variations in absolute fluorescence values. Results are presented as representative outcomes of experiments conducted at least three times independently.

### 9. PEG switch off assay

TCA-based denatured extracts were resuspended in PB buffer (8 M urea, 0.2 M pH 8.5 Tris, 0.5% SDS, 0.1% deoxycholic acid, 0.5% Triton-X-100) for 30 minutes. To block non-oxidized cysteine residues, 0.1 M iodoacetamide (IAM) was added, and the mixture was incubated for 1 hour at room temperature in darkness. Unbound IAM was removed by TCA precipitation (30 minutes at -20°C followed by centrifugation at 4°C and 21,300 x g for 10 minutes). Subsequently, oxidized cysteine residues were resuspended in PB buffer supplemented with tris(2-carboxyethyl) phosphine hydrochloride (TCEP) and incubated for 30 minutes at room temperature. Methoxy PEG-maleimide was added directly to the buffer to achieve a final concentration of 3 mg/ml and incubated for an additional one hour at room temperature. The mixture was then subjected to TCA precipitation to remove unbound Methoxy PEG-maleimide. Finally, the pellet was resuspended in non-reduced Laemmli sample buffer for subsequent immunoblotting.

### 10. Drop assay for cells survival following acute oxidative stress

Overnight yeast cultures were diluted to 0.5 OD600/ml in YP medium supplemented with 4% glucose. Cultures were then incubated at 28°C for 4 hours before being treated with high concentrations of 0-40 mM H_2_O_2_ for 1 hour. The survival of mutants and WT cells was assessed using drop assays. Treated cell cultures were serially diluted 1:5 in YP medium before spotting on YPAD or YPGly plates (YP with 3% glycerol). The plates were incubated at 28°C for 2-4 days, after which cell viability was determined.

### 11. Fluorescence Microscopy

Overnight cultures of yeast strains expressing IDP-GFP were diluted to 0.5 OD600 in SC medium selective to uracil, supplemented with 4% glucose. Cultures were then incubated at 28°C for 4 hours. Cell morphology, as well as differential interference contrast (DIC) and fluorescence images of cells, were captured using an Olympus BX53 Digital Fluorescence Microscope (EVIDENT CORPORATION, Tokyo, Japan). The morphology of all mitochondria observed in cells was assessed based on ≥ 5 independent photographs, as previously described ^46^.

### 12. Mitochondria volume

Mitochondria of cells expressing IDP1-GFP were observed by Nikon A1R Confocal Laser Scanning Microscope at the Bioimaging Unit of the Faculty of Natural Sciences - University of Haifa. Imaris 9.8 (Oxford Instruments) was used for surface rendering using the “Surfaces” module to determine the object volumes. For quantification of mitochondrial volumes, thresholds for surface creation were guided by automatic thresholding.

### 13. Statistical analyses

One-way ANOVA was applied to study the differences between the mitochondrial volume to cell-area ratios of WT, *Δrub1* and *Δyuh1*. Two-way ANOVA was applied to study the differences between the mitochondrial morphotypes (tubular vs. non-tubular) of the three strains. All statistical analyses were performed on IBM SPSS 25 software. All groups were normally distributed according to Shapiro-Wilk tests (p>0.05). The error variances were equal across the three groups according to Leven’s tests based on both means and medians (p>0.05).

## Bibliography

1. Barford, D. The role of cysteine residues as redox-sensitive regulatory switches. Curr Opin Struct Biol 14, 679–86 (2004).

2. Schilter, D. Thiol oxidation: A slippery slope. Nature Reviews Chemistry 1, 0013 (2017).

3. Stankovic-Valentin, N. & Melchior, F. Control of SUMO and Ubiquitin by ROS: Signaling and disease implications. Mol Aspects Med (2018).

4. Cappadocia, L. & Lima, C.D. Ubiquitin-like Protein Conjugation: Structures, Chemistry, and Mechanism. Chem Rev (2017).

5. Tolbert, B.S. et al. The active site cysteine of ubiquitin-conjugating enzymes has a significantly elevated pKa: functional implications. Biochemistry 44, 16385–91 (2005).

6. Jahngen-Hodge, J. et al. Regulation of ubiquitin-conjugating enzymes by glutathione following oxidative stress. J Biol Chem 272, 28218–26 (1997).

7. Yao, D. et al. Nitrosative stress linked to sporadic Parkinson’s disease: S-nitrosylation of parkin regulates its E3 ubiquitin ligase activity. Proc Natl Acad Sci U S A 101, 10810–4 (2004).

8. Whitby, F.G., Xia, G., Pickart, C.M. & Hill, C.P. Crystal Structure of the Human Ubiquitin-like Protein NEDD8 and Interactions with Ubiquitin Pathway Enzymes. J. Biol. Chem. 273, 34983–34991 (1998).

9. Soucy, T.A. et al. An inhibitor of NEDD8-activating enzyme as a new approach to treat cancer. Nature 458, 732–6 (2009).

10. Mahon, C., Krogan, N.J., Craik, C.S. & Pick, E. Cullin E3 ligases and their rewiring by viral factors. Biomolecules 4, 897–930 (2014).

11. Sarikas, A., Hartmann, T. & Pan, Z.Q. The cullin protein family. Genome Biol 12, 220 (2011).

12. Shen, L.N. et al. Structural basis of NEDD8 ubiquitin discrimination by the deNEDDylating enzyme NEDP1. EMBO J 24, 1341–51 (2005).

13. Wada, H., Kito, K., Caskey, L.S., Yeh, E.T. & Kamitani, T. Cleavage of the C-terminus of NEDD8 by UCH-L3. Biochem Biophys Res Commun 251, 688–92 (1998).

14. Schmaler, T. & Dubiel, W. Control of Deneddylation by the COP9 Signalosome. Subcell Biochem 54, 57–68 (2010).

15. Dubiel, D., Rockel, B., Naumann, M. & Dubiel, W. Diversity of COP9 signalosome structures and functional consequences. FEBS Lett 589, 2507–13 (2015).

16. Bramasole, L. et al. Proteasome lid bridges mitochondrial stress with Cdc53/Cullin1 NEDDylation status. Redox Biol 20, 533–543 (2019).

17. Kumar, A. et al. The bacterial fermentation product butyrate influences epithelial signaling via reactive oxygen species-mediated changes in cullin-1 neddylation. J Immunol 182, 538–46 (2009).

18. Reisz, J.A., Bechtold, E., King, S.B., Poole, L.B. & Furdui, C.M. Thiol-blocking electrophiles interfere with labeling and detection of protein sulfenic acids. Febs j 280, 6150–61 (2013).

19. Bossis, G. & Melchior, F. Regulation of SUMOylation by reversible oxidation of SUMO conjugating enzymes. Mol Cell 21, 349–57 (2006).

20. Saad, S., Berda, E., Klein, Y., Issa, S. & Pick, E. Strategies for Monitoring “Ubiquitin C-Terminal Hydrolase 1” (Yuh1) Activity. Methods Mol Biol 2602, 107–122 (2023).

21. Kohr, M.J. et al. Simultaneous measurement of protein oxidation and S-nitrosylation during preconditioning and ischemia/reperfusion injury with resin-assisted capture. Circ Res 108, 418–26 (2011).

22. Harshuk-Shabso, D., Castel, N., Israeli, R., Harari, S. & Pick, E. Saccharomyces cerevisiae as a Toolkit for COP9 Signalosome Research. Biomolecules 11(2021).

23. Gurevich, S.Z. et al. Rub1/NEDD8, a ubiquitin-like modifier, is also a ubiquitin modifier. bioRxiv, 2020.06.18.159145 (2020).

24. Starkov, A.A. The role of mitochondria in reactive oxygen species metabolism and signaling. Ann N Y Acad Sci 1147, 37–52 (2008).

25. Egner, A., Jakobs, S. & Hell, S.W. Fast 100-nm resolution three-dimensional microscope reveals structural plasticity of mitochondria in live yeast. Proc Natl Acad Sci U S A 99, 3370–5 (2002).

26. Tsuboi, T. et al. Mitochondrial volume fraction and translation duration impact mitochondrial mRNA localization and protein synthesis. Elife 9(2020).

27. Hoffmann, H.P. & Avers, C.J. Mitochondrion of yeast: ultrastructural evidence for one giant, branched organelle per cell. Science 181, 749–51 (1973).

28. Stevens, B.J. & White, J.G. Computer reconstruction of mitochondria from yeast. Methods Enzymol 56, 718–28 (1979).

29. Kondo, W., Kitagawa, T., Hoshida, H., Akada, R. & Miyakawa, I. Morphological Changes of Mitochondria and Actin Cytoskeleton in the Yeast <i>Saccharomyces cerevisiae</i> During Diauxic Growth and Glucose Depletion Culture. CYTOLOGIA 87, 157–162 (2022).

30. Chelius, X. et al. Selective retention of dysfunctional mitochondria during asymmetric cell division in yeast. PLoS Biol 21, e3002310 (2023).

31. Abeliovich, H., Zarei, M., Rigbolt, K.T., Youle, R.J. & Dengjel, J. Involvement of mitochondrial dynamics in the segregation of mitochondrial matrix proteins during stationary phase mitophagy. Nat Commun 4, 2789 (2013).

32. Journo, D., Mor, A. & Abeliovich, H. Aup1-mediated regulation of Rtg3 during mitophagy. J Biol Chem 284, 35885–95 (2009).

33. Ben-Lulu, S., Ziv, T., Admon, A., Weisman-Shomer, P. & Benhar, M. A substrate trapping approach identifies proteins regulated by reversible S-nitrosylation. Mol Cell Proteomics 13, 2573–83 (2014).

34. Keuss, M.J. et al. Unanchored tri-NEDD8 inhibits PARP-1 to protect from oxidative stress-induced cell death. EMBO J 38(2019).

35. Goellner, E.M. et al. PCNA and Msh2-Msh6 activate an Mlh1-Pms1 endonuclease pathway required for Exo1-independent mismatch repair. Mol Cell 55, 291–304 (2014).

36. Costanzo, M. et al. The genetic landscape of a cell. Science 327, 425–31 (2010).

37. Costanzo, M. et al. A global genetic interaction network maps a wiring diagram of cellular function. Science 353(2016).

38. Downs, C.A., Kumar, A., Kreiner, L.H., Johnson, N.M. & Helms, M.N. H2O2 regulates lung epithelial sodium channel (ENaC) via ubiquitin-like protein Nedd8. J Biol Chem 288, 8136–45 (2013).

39. Collier-Hyams, L.S., Sloane, V., Batten, B.C. & Neish, A.S. Cutting edge: bacterial modulation of epithelial signaling via changes in neddylation of cullin-1. J Immunol 175, 4194–8 (2005).

40. Pajares, M. et al. Redox control of protein degradation. Redox Biol 6, 409–420 (2015).

41. Lefaki, M., Papaevgeniou, N. & Chondrogianni, N. Redox regulation of proteasome function. Redox Biol 13, 452–458 (2017).

42. Manford, A.G. et al. A Cellular Mechanism to Detect and Alleviate Reductive Stress. Cell (2020).

43. Silva, G.M., Finley, D. & Vogel, C. K63 polyubiquitination is a new modulator of the oxidative stress response. Nat Struct Mol Biol 22, 116–23 (2015).

44. Aiken, C.T., Kaake, R.M., Wang, X. & Huang, L. Oxidative stress-mediated regulation of proteasome complexes. Mol Cell Proteomics 10, R110.006924 (2011).

45. Bailly, A. et al. The NEDD8 inhibitor MLN4924 increases the size of the nucleolus and activates p53 through the ribosomal-Mdm2 pathway. Oncogene (2015).

46. Sinha, A. et al. The COP9 signalosome mediates the Spt23 regulated fatty acid desaturation and ergosterol biosynthesis. FASEB J (2020).

47. Yu, Z. et al. Dual function of Rpn5 in two PCI complexes, the 26S proteasome and COP9 signalosome. Mol Biol Cell 22, 911–20 (2011).

