## Supplemental Data for "Redox-driven control of Yuh1/UCHL3 impacts mitochondrial health via NEDD8/Rub1 pathway"

Address

**On the following pages, this file contains the following sections:**

A) **7 supplementary Figures**
B) **4 Supplementary Tables (3 enclosed and one excel doc')**
C) **Supplementary materials and methods**
D) **Supplementary references**

#### A) SUPPLEMENTARY FIGURES (1-7):

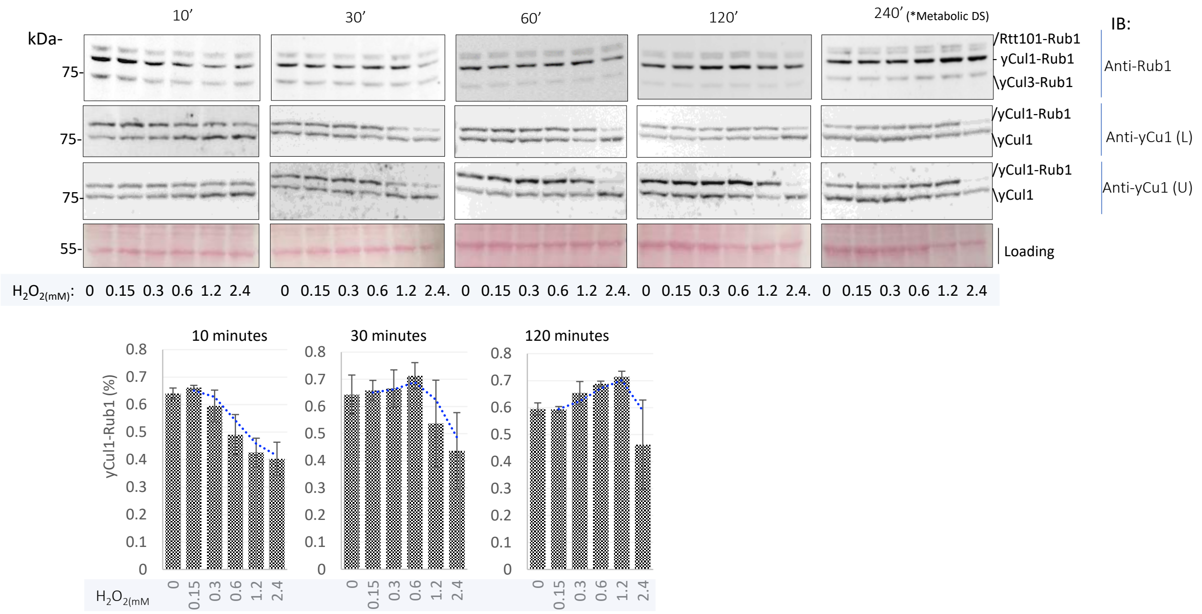


**Figure S1: Dynamic assessment of cullin NEDDylation in response to time-dependent $H_2O_2$-induced reactive oxygen species (ROS).** Logarithmic-phase-grown cultures of wild type cells expressing genome-engineered *UBC12* ORF, tagged with 3HA were subjected to a change in growth medium, replacing it with an equal volume of fresh SD medium supplemented with 0-2.4 mM $H_2O_2$ over time. Importantly, in 240 minutes, cultures reached the endogenous metabolic diauxic shift (DS). Cells were harvested for immunoblotting to detect cullins by antibodies for Rub1 (recognizing the three cullins), or two independent antibodies developed for yCul1, one of them preferentially detects the NEDDylated form (U), while the other one targets the free form (L). Quantification of yCul1 NEDDylation status was carried out using ImageJ software ($n \geq 3$).

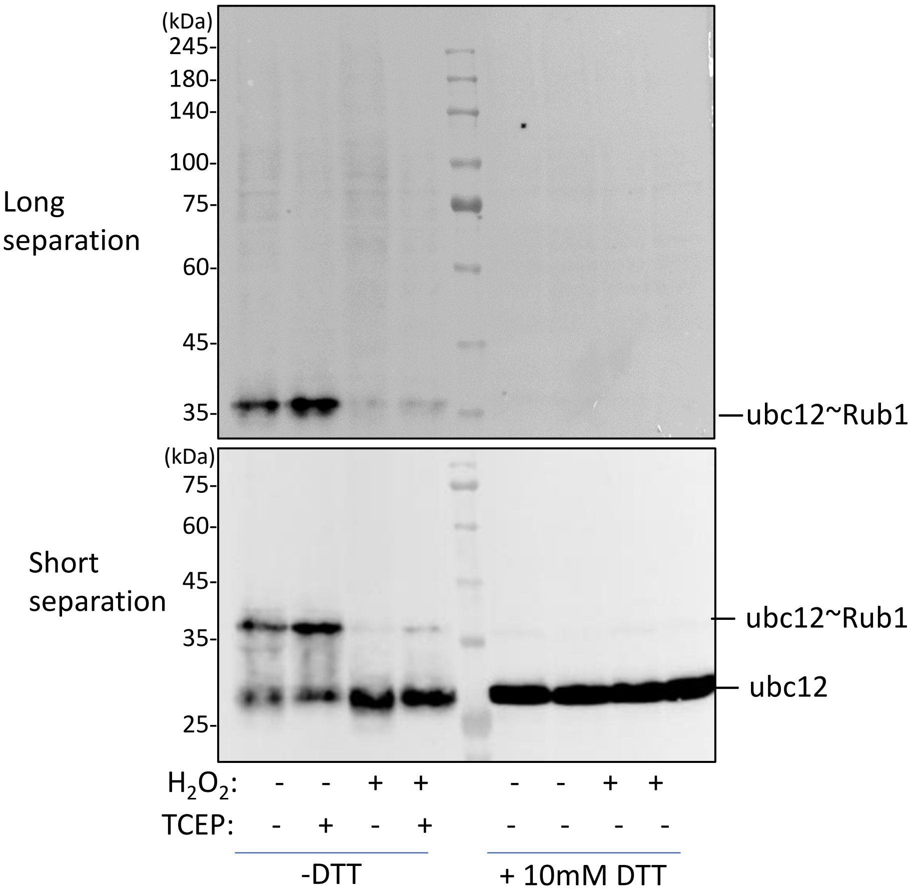


**Figure S2: Changes in Ubc12~Rub1 thioester forms following $H_2O_2$ treatment.** Logarithmic phase WT cells expressing endogenous Ubc12-HA were treated with 2.4 mM $H_2O_2$ and incubated for 10 minutes. High molecular weight forms (above 35 kDa) were examined using immunoblotting with non-reducing urea sample buffer, 50mM TCEP urea sample buffer, or 100mM DTT urea sample buffer.

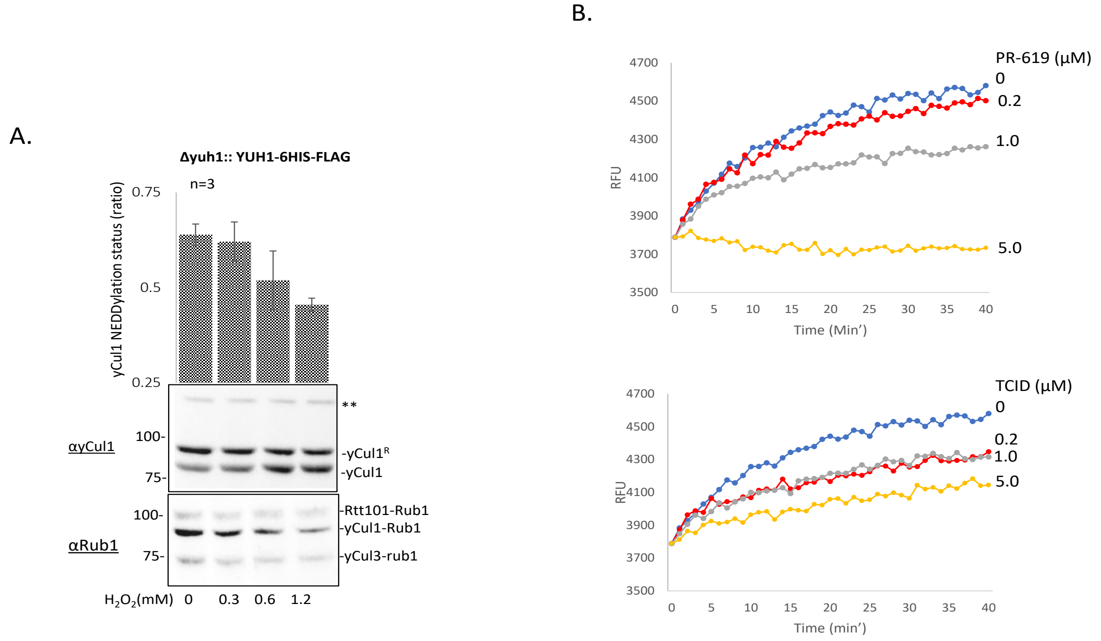


**Figure S3: Inhibition of Yuh1 Activity by PR619 and TCID.** Δ*yuh1* mutant strain complemented by 6His-Flag-YUH1 under *YUH1p* (Yuh1-WT) was grown to the logarithmic phase at 28°C, and native lysates were prepared. (A) The lysates were subjected to immunoblotting for yCul1 and anti-Rub1, which recognizes the three cullins. The detection of the neddylation cascade in this transformant indicates its sensitivity to oxidative stress, thus validating the strain for further experiments. A nonspecific band was observed (**), serving as the loading control. (B) The lysates were also used to assess Yuh1 activity by measuring the ability to cleave the amide bond between Gly76 of ubiquitin and 7-amido-4-methylcoumarin (AMC), with or without the addition of various doses of the TCID (Cayman 30675) and PR619 (Cayman 2645-32-1). This was quantified in relative fluorescence units (RFU) using a plate reader (excitation 360/40 nm and emission 460/40 nm), illustrating notable inhibition of Yuh1 activity.

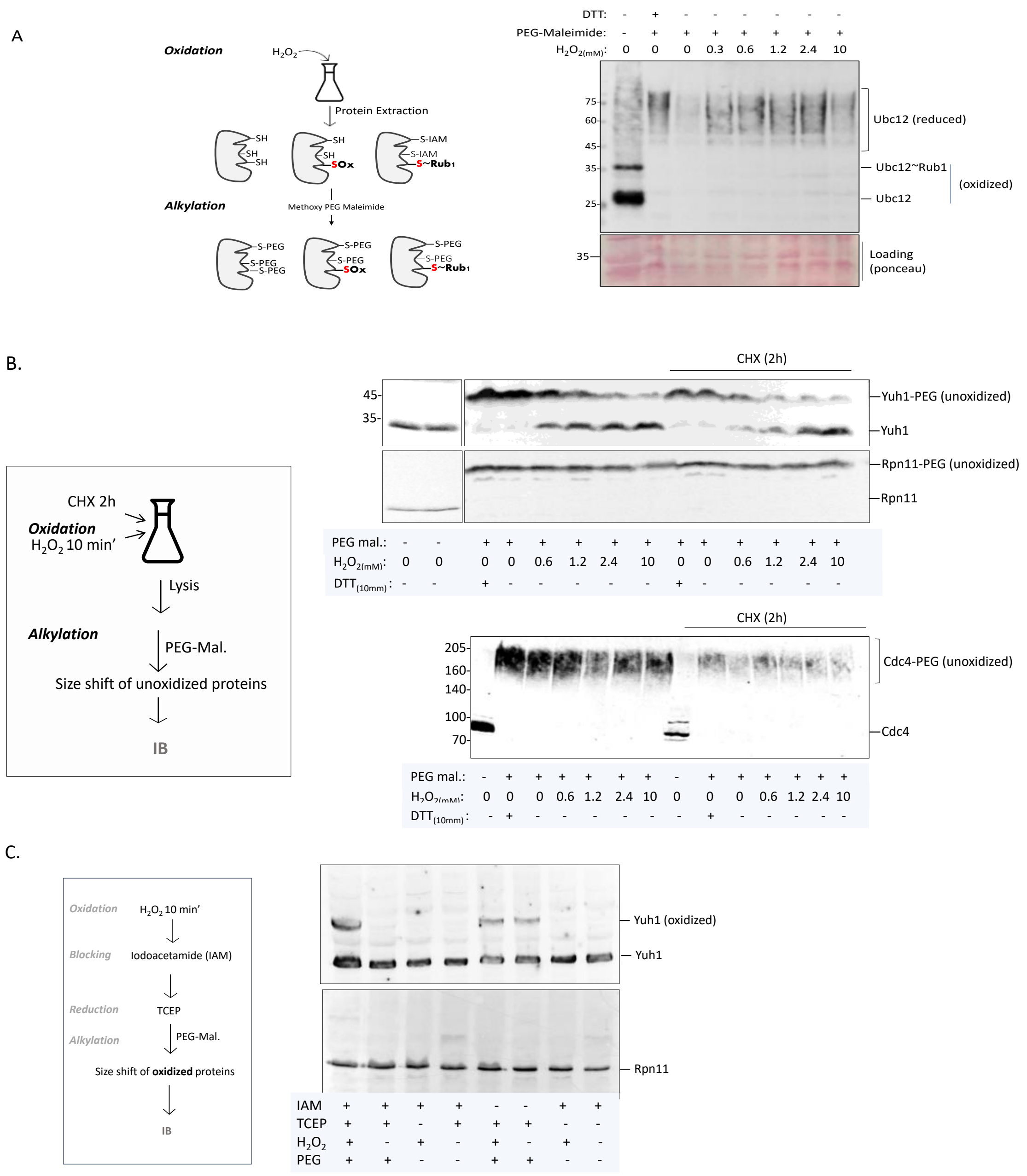


**Figure S4: Searching for oxidized Cys residues in the NEDDylation cascade through gel shift assays.** Logarithmic Δ*yuh1* mutant cells, co-expressing endogenous Ubc12-HA and ectopic Yuh1-6His-Flag were treated with the indicated concentrations of $H_2O_2$ for 10 minutes or 10 mM DTT for 30 minutes. (A) To monitor changes in reduced Cys, 3 mg/ml methoxy-polyethylene glycol maleimide (PEG-Mal) was added, as illustrated in the scheme (left). Total protein samples were separated by non-reducing protein SDS-PAGE and subjected to immunoblotting. (B) The same experimental setup was followed a two-hour treatment of yeast cells with cycloheximide (CHX) to inhibit protein expression, allowing observation only after protein degradation. Rpn11 served as a control for a stable protein that is not sensitive to oxidation, possessing a single cysteine. Cdc4 was used as a control for a short-lived protein, confirming the effectiveness of CHX. (C) Following extraction, a PEG switch off assay was performed as indicated and anti-Flag antibodies were used to recognize the shift size of Yuh1.

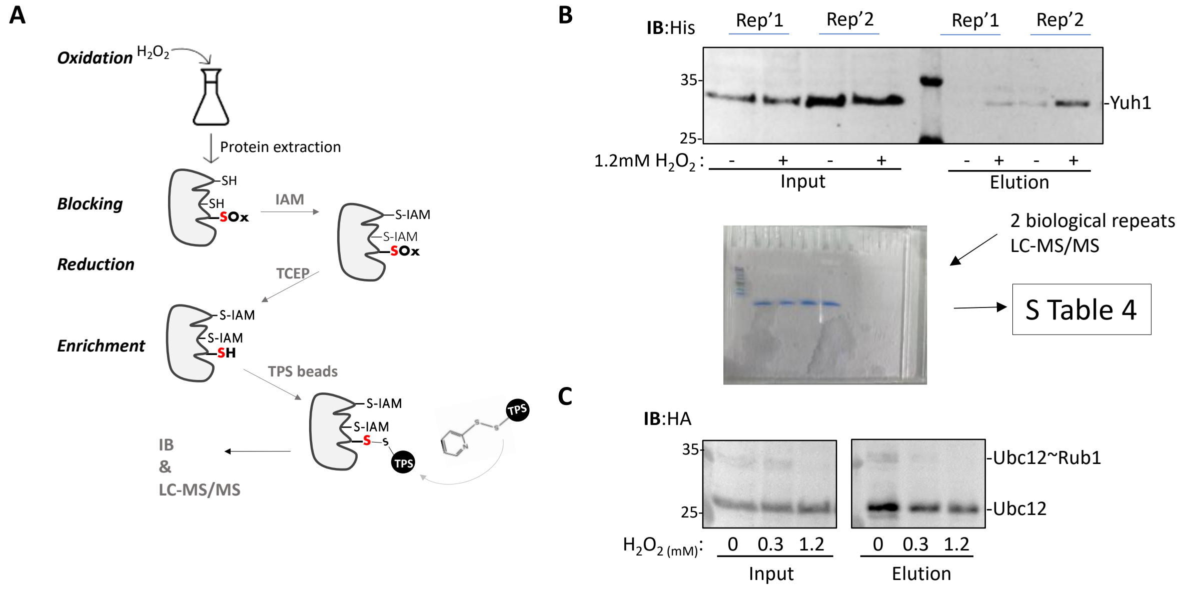


**Figure S5: Assessment of Ubc12 Oxidation using Resin-Assisted Capture (Ox-RAC) Assay.**

(A) A schematic representation of the experiment wherein free thiols are blocked by Iodoacetamide (IAM), and oxidized thiols are reduced by TCEP. Following reduction, these thiols can form disulfides with the TPS beads before elution under reducing conditions by TCEP. (B) Logarithmic Δ*yuh1* mutant cells, co-expressing endogenous Ubc12-HA and ectopic Yuh1-6His-Flag were treated with indicated concentrations of $H_2O_2$ for 10 minutes. Ox-RAC was carried out using 100 mM iodoacetamide (IAM) as a blocker and 10 mM TCEP to reduce the previously oxidized Cys prior to the binding to TPS beads and elution with 50 mM TCEP. Yuh1 (B) and (C) Ubc12 levels in the pulldown were examined by immunoblotting for Ubc12-HA. Samples were also separated halfway in an SDS PAGE (B, bottom), and bands were cut out of the gel for protein extraction. The extracted proteins were trypsinized, and the peptides were analyzed by LC-MS/MS. The complete dataset is available in an Excel sheet provided in Table S4.

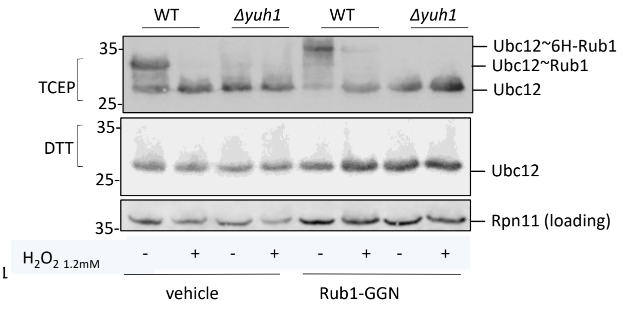


**Figure S6: Yuh1 is Essential for Rub1 Incorporation into the NEDDylation Cascade.** Logarithmic cultures of WT Ubc12-HA and Δ*yuh1* Ubc12-HA mutant strains expressing the Rub1 precursor (Rub1-GGN) were treated with1.2mM of $H_2O_2$ for 10 minutes and subsequently harvested for immunoblotting under non-reduced (top) or reduced (bottom) conditions. The results indicate that in the absence of Yuh1, Rub-GGN remains unprocessed, and cullins are not NEDDylated.

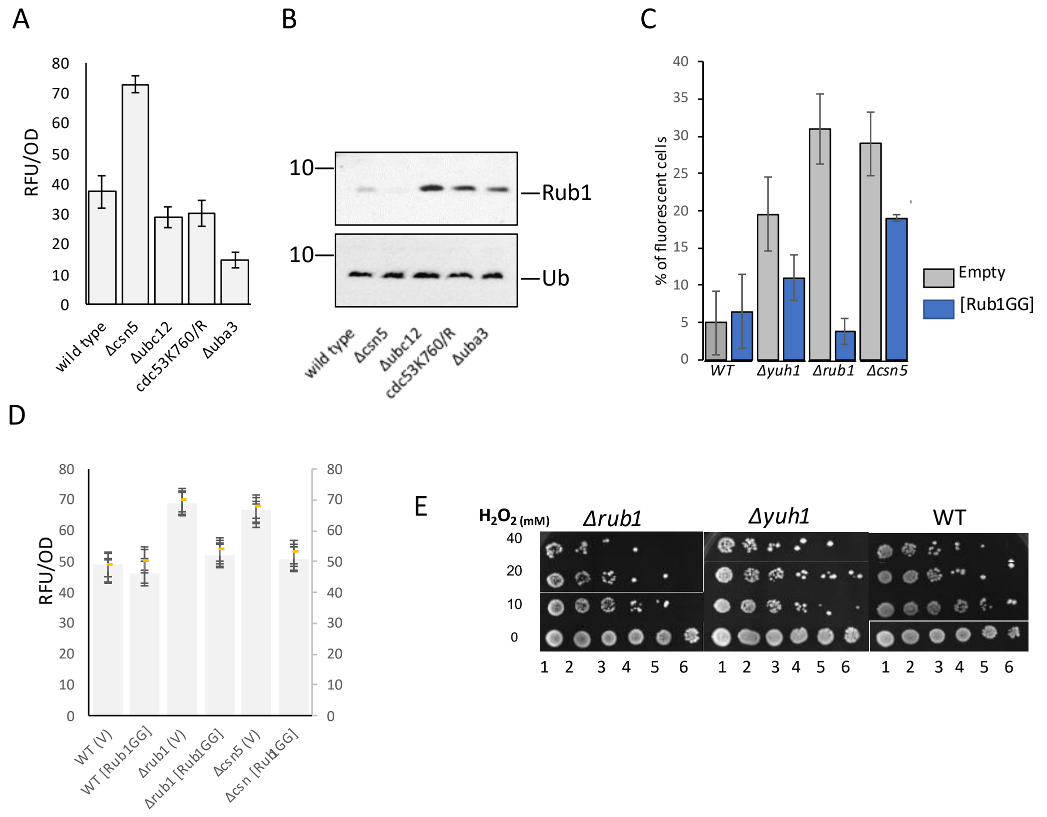

**Figure S7: Oxidation state in strains with varied free Rub1 levels.** (A, B) WT yeast strain, along with the *Δcsn5* mutant strain characterized by reduced levels of free Rub1, and strains lacking the NEDDylation components *Δubc12*, *Δuba3*, and *cdc53 K760/R*, which are characterized by accumulated non-NEDDylated cullins and elevated free Rub1 levels, were cultured at 28°C. Intracellular ROS levels were quantified using a BioTek Synergy HT plate reader (Excitation=485/20; Emission=528/20) (A). (B) Protein extracts were subjected to immunoblotting for Rub1 and ubiquitin. (C) WT, Δ*yuh1*, Δ*rub1* and Δ*csn5* mutant strains expressing empty vector or Rub1-GG were cultures until the diauxic shift (12 hours). One hour prior to harvest, cells were treated with 2',7' –dichlorofluorescin diacetate (DCFDA) to evaluate ROS levels using a plate reader, as described. (D) Rub1GG was overexpressed in strains with low Rub1 levels, with vector (V) used as a control, and intracellular ROS levels were measured using a plate reader as described above. (E) Logarithmically growing WT, Δ*yuh1*, Δ*rub1* and Δ*csn5* mutant cells were treated with 0-40 mM $H_2O_2$. Recovery was assessed by seeding treated cultures onto YPAD plates using a serial dilution assay.

### SUPPLEMENTARY TABLES (1-3)

S Table 1: List of plasmids used in this study

| **Plasmid** | **Characteristics** | **Source** |
|---|---|---|
| EP53 | Yeplac181 empty plasmid, 2µ plasmid, *ADH*, amp/leu | Open Biosystems |
| EP115 | RGS-Rub1ΔN77(intron free) plasmid, 2µ plasmid, *ADH*, amp/leu | Open Biosystems |
| EP181 | RGS-Rub1(intron-free) + 3' UTR plasmid, 2µ plasmid, *ADH*, amp/leu | Open Biosystems |
| EP184 | Yeplac181 empty plasmid, 2µ plasmid, amp/leu | Sugino |
| EP278 | *YUH1p*-YUH1-6His-Flag in Yeplac181 plasmid, 2µ, amp/leu | This study |
| EP291 | RGS-6(HIS)-UBB+1 in pQE30 plasmid, T5, amp | Open Biosystems |
| EP292 | 6His-Ulp1 in pET28b plasmid, T7, kan | [1] |
| EP294 | Yuh1 in pET3a plasmid, T7, amp | [2] |
| EP110 | IDP1-GFP, amp/ura | Open Biosystems |
| EP287 | 6His-Ulp1 in pET28b, T7, Kan | [1] |

S Table 2: list of yeast strains used in this study

| **Strain** | **Mat** | **Genotype** | **Source** |
|---|---|---|---|
| YP146 | *α* | *ura3Δ0 leu2Δ0 his3Δ1; Δcsn5 YDL216c:: KanMX4* | EUROSCARF |
| YP376 | a | *ura3Δ0 leu2Δ0 his3Δ1 met15Δ0; CDC53::KanMX4 CDC53(k760r)::LEU* | Open Biosystems |
| YP396 | a | WT *ura3Δ0 leu2Δ0 his3Δ1 met15Δ0* | Open Biosystems |
| YP468 | *α* | *ura3Δ0 leu2Δ0 his3Δ1 lys2Δ0; UBC12::UBC12-HA-HIS3* | [3] |
| YP484 | *a* | *ura3Δ0 leu2Δ0 his3Δ1; Δubc12 YLR306W:: KanMX4* | EUROSCARF |
| YP487 | a | *ura3Δ0 leu2Δ0 his3Δ1 met15Δ0; Δuba3 YPR066w::KanMX4* | EUROSCARF |
| YP540 | a | *ura3Δ0 leu2Δ0 his3Δ1; met15Δ0; rub1::G418* | This study |
| YP545 | *a* | *ura3Δ0 leu2Δ0 his3Δ1; lys2Δ0 met15Δ0; yuh1:: KanMX4* | This study |
| YP546 | a | *ura3Δ0 leu2Δ0 his3Δ1 lys2Δ0; UBC12:: UBC12-HA-HIS3; Yuh1:: KanMX4* | This study |

S Table 3: list of antibodies used in this study

| Primary | Secondary | Source | Catalog no. | Primary dilution |
|---|---|---|---|---|
| Cdc53 (Cull) | Rabbit | Santa Cruz | Sc 50444 | 1:1000 |
| Cdc53 (Cull) | Goat | Santa Cruz | Sc 6717 | 1:1000 |
| Rpn11 | Rabbit | [4] Yu et al. 2011 | | 1:1000 |
| Hemagglutinin (HA) | Mouse | Santa Cruz | Sc 7392 | 1:1000 |
| Rub1 | Rabbit | Abcam | ab 4751sc | 1:500 |
| His tag | Mouse | GeneScript | A00186 | 1:1000 |
| Cdc4 | Goat | Santa Cruz | Sc 6715 | 1:1000 |
| Ubiquitin | Rabbit | Dako | Z0458 | 1:500 |
| Flag tag | Mouse | Genscript | A00187 | 1:1000 |

1. Supplementary information for S Table 4 (Excel sheet)

The proteins separated by SDS-PAGE were subjected to trypsin digestion, and the resulting peptides were analyzed by LC-MS/MS using a Q Exactive HFX mass spectrometer (Thermo). The data were processed with MaxQuant 1.5.2.8, with a false discovery rate (FDR) cutoff of $<0.01$ applied. Contaminating proteins were removed from the dataset. Protein groups were compiled to present data at the protein level, incorporating all evidence for each specific protein. A ratio was calculated between the control and experimental conditions, with specific proteins highlighted in orange. Missing values in each report were replaced with 100,000, serving as a threshold level.

2. Oxidized cysteine resin-assisted capture method (OxRAC)

TCA pellets of $H_2O_2$ treated cells were carefully resuspended using urea buffer pH 9, consisting of 8M urea and 230mM Tris pH 11 in HEN buffer (comprising 250mM HEPES, 1mM EDTA, and 0.1M neocuproine adjusted to pH 7.5). Ensuring equal protein concentrations were maintained, the next step involved blocking non-oxidized cysteine thiols with 100mM iodoacetamide (IAM) at room temperature for 1 hour in darkness. Following thiol blocking, acetone precipitation was performed to eliminate unbound IAM. This involved adding 100% acetone to the samples and incubating them at -20°C for 30 minutes. Subsequent centrifugation at 13,000g for 30 minutes at 4°C facilitated the collection of protein pellets. To reverse the oxidation of cysteine residues, the protein pellets were resuspended in a buffer consisting of 10mM TCEP pH 7, 8M urea, 230mM Tris pH 6.8, and 1% SDS in HEN buffer. Incubation at room temperature in darkness for 1 hour allowed for effective reduction of oxidized cysteines. Excess reducing agent was removed through a series of washes with 70% acetone, followed by centrifugation at 13,000g for 5 minutes at 4°C after each wash. The final pellet was then resuspended in 1% SDS HEN (HENS) buffer and pre-washed thiopropyl-sepharose (TPS) beads were added. For binding, tubes were incubated overnight with rotation in the dark at 4°C, facilitating the binding of proteins to the TPS beads. The TPS-bound proteins were then washed four times with HENS buffer, followed by two washes with 10x diluted HENS buffer (10% HENS). Elution from the beads was carried out by elution buffer (50 mM TCEP in 10% HENS), and used for downstream analyses of immunoblotting and MS.

### SUPPLEMENTARY REFERENCES

1. Catanzariti A, Soboleva TA, Jans DA, Board PG, Baker RT. An efficient system for high-level expression and easy purification of authentic recombinant proteins. *Protein Science* **13(5)**,1331-1339 (2004).

2. Johnston SC, Riddle SM, Cohen RE, Hill CP. Structural Basis for the Specificity of Ubiquitin C-Terminal Hydrolases. *The EMBO Journal* **18**, 3877-3887 (1999).

3. Rabut, Gwenaël et al. The TFIIH subunit Tfb3 regulates cullin neddylation. *Molecular cell* **43(3)**, 488–95 (2011).

4. Yu Z, Kleifeld O, Lande-Atir A, et al. Dual function of Rpn5 in two PCI complexes, the 26S proteasome and COP9 signalosome. *Molecular biology of the cell* **22(7)**, 911-20 (2011).
